# The C. elegans Notch proteins LIN-12 and GLP-1 are tuned to lower force thresholds for activation than Drosophila Notch

**DOI:** 10.1101/2021.02.11.429991

**Authors:** Paul D. Langridge, Jessica Yu Chan, Alejandro Garcia-Diaz, Iva Greenwald, Gary Struhl

## Abstract

The conserved transmembrane receptor Notch mediates cell fate decisions in all animals. In the absence of ligand, a Negative Regulatory Region (NRR) in the Notch ectodomain adopts an autoinhibited confirmation, masking an ADAM protease cleavage site [1, 2]; ligand binding makes the cleavage site accessible, leading to shedding of the Notch ectodomain as the first step of signal transduction [3, 4]. In Drosophila and vertebrates, the ligands are all single-pass transmembrane Delta/Serrate/LAG-2 (DSL) proteins; the endocytic adaptor Epsin binds to the ubiquitinated intracellular domain, and the resulting Clathrin-mediated endocytosis exerts a “pulling force” that exposes the cleavage site in the NRR [4–6]. However, in *C. elegans*, the presence of natural secreted DSL proteins [7] and other observations suggested that Epsin-mediated endocytosis may not be required to activate the Notch proteins LIN-12 and GLP-1. Here, we confirm that neither Epsin nor the cytosolic domains of DSL proteins are required for Notch signaling in C. elegans. Furthermore, we provide evidence that the NRRs of LIN-12 and GLP-1 are tuned to a lower force level than the NRR of Drosophila Notch. Finally, we show that adding a Leucine “plug” that occludes the cleavage site in vertebrate and Drosophila Notch proteins but is absent in the *C. elegans* Notch proteins [1, 2] renders the LIN-12 and GLP-1 NRRs dependent on Epsin-mediated ligand endocytosis, indicating that greater force is now required to expose the cleavage site. Thus, the NRRs of LIN-12 and GLP-1 appear to be tuned to a lower force threshold, accounting for the different requirements for signaling in *C. elegans.*

## Results

### Neither Epsin nor direct interactions with other cytosolic proteins are required for DSL ligands to signal in *C. elegans*

Epsin-mediated endocytosis of ligand is an obligate requirement for DSL/Notch signaling in Drosophila and mammals: when Epsin activity is removed from signaling cells, they are unable to activate Notch in receiving cells, resulting in phenotypes that are indistinguishable from those caused by Notch loss-of-function mutations [8–12]. *C. elegans* has a single Epsin ortholog, EPN-1 [10, 13, 14], and *epn-1* null mutants *[epn-1(0)]* arrest during embryogenesis [14]. As the absence of *Notch* signaling also results in embryonic lethality, we began by investigating whether the *epn-1* null mutants arrest with cell fate transformations resulting from a failure in DSL/Notch signaling.

Loss of DSL/Notch signaling is lethal because of the failure of two ligands LAG-2 and APX-1 to execute their essential roles in cell-fate specification. To investigate the potential requirement of Epsin for ligand function, we examined *epn-1(0)* mutants for defects associated with the absence of zygotic *lag-2* activity [15] or maternal *apx-1* activity [16].

Loss of *lag-2* results in fully penetrant lethality [15], with hallmark cell fate transformations that are also seen in the *lin-12 glp-1* double mutant because the two *C. elegans* Notch proteins, LIN-12 and GLP-1, are functionally redundant for certain cell-cell interactions during embryogenesis [15]. The most striking defects are the absence of an excretory cell and the absence of a rectum, which result because of a failure of inductive signaling from MS descendants to ABp descendants [17]. We generated *epn-1(0)* individuals by loss of a “simple” extrachromosomal array that appears to have little or no maternal contribution [14] and observed that all homozygous *epn-1(0)* mutants have an excretory cell, which could be readily visualized with a fluorescent marker (Figure 1A); *epn-1(0)* mutants also had a rectum (data not shown). These results indicate that *epn-1* is not required for LAG-2 to activate LIN-12 and/or GLP-1 during zygotic events that require LAG-2 activity, and is further supported by mutational analysis of LAG-2 below.

**Figure 1.**
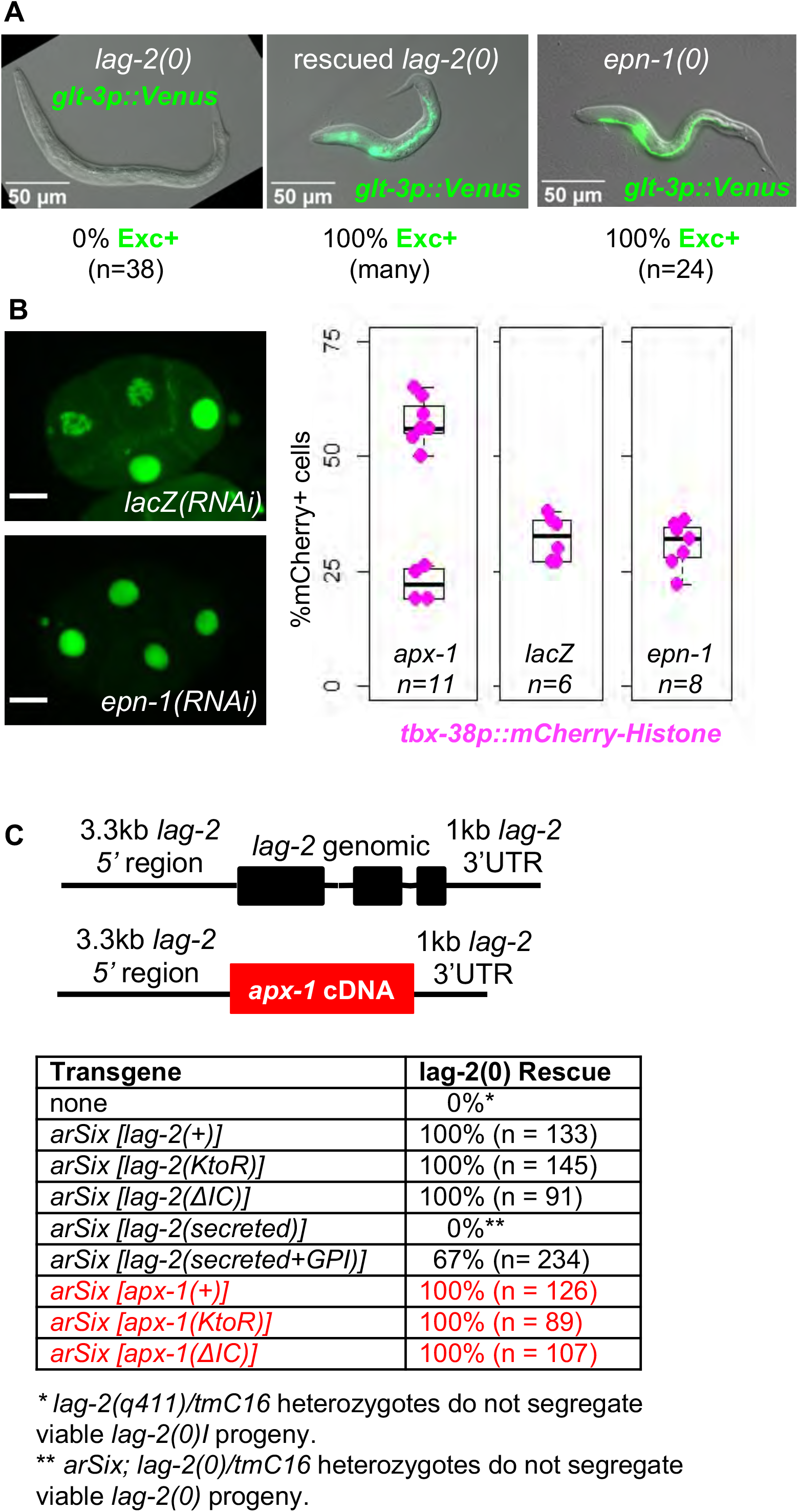
Epsin is not required for DSL protein ligand activity in *C. elegans*. **A. *epn-1(0)* mutants do not display defects associated with the absence of zygotic *lag-2* activity**. The excretory cell is specifically labeled by *arIs164[glt-3p::Venus]* [48]. A *lag-2(0)* mutant (left) lacks an excretory cell; such mutants are obtained from a rescued strain carrying an extrachromosomal array, and rescued individuals have an excretory cell (middle). Homozygous *epn-1(0)* mutants may also be obtained as segregants from a rescued strain carrying a”simple” extrachromosomal array that appears to have little or no maternal contribution [14], and such *epn-1(0)* homozygotes always have an excretory cell (right). **B. *epn-1(0)* mutants do not display excess pharyngeal cells associated with the absence of maternal *apx-1* activity**. We used RNAi to deplete *epn-1* activity in early embryos by feeding parents bacteria expressing double-stranded RNA (STAR Methods). HIS-72::GFP was used to mark all nuclei at all stages, and mCherry:HistoneH1 to mark pharyngeal cells to facilitate scoring cell fate [19]. Endogenously-tagged EPN-1::GFP is visible in the cytoplasm and somewhat enriched at cell membranes (left, top) in *lacZ(RNAi)* embryos; in the experiment shown, we identified *epn-1(RNAi)* embryos that lacked detectable EPN-1::GFP at the four-cell stage (left, bottom), and observed the same embryos again later for the proportion of pharyngeal cells that formed. Normal late-stage embryos have about 25% pharyngeal cells whereas loss of *apx-1* activity results in about 50% pharyngeal cells; the difference between *apx-1(RNAi)* and *epn-1(RNAi)* was significant at p<0.05 (Fisher’s exact test). We also performed a similar experiment without following specific embryos; 12/12 *epn-1(RNAi)* individuals had normal numbers of pharyngeal cells whereas 14/22 *apx-1(RNAi)* embryos had excess pharyngeal cells; this difference was significant at p<0.01 (Fisher’s exact test). We note that in all cases, the *epn-1(RNAi)* was effective enough to result in inviable embryos. **C. *lag-2(0)* rescue experiments**. Schematic drawings (not to scale) of constructs inserted into a defined site on LGI and assayed for rescue of the lethality caused by a *lag-2* null allele (see STAR Methods). In addition to rescue of lethality shown, *arIs51[cdh-3p::GFP]* was used to score the number of ACs when the intracellular domain was deleted: 19/19 individuals of genotype *lag-2(ΔIC); lag-2(0); arIs51[cdh-3p::GFP]* and 18/18 *apx-1(ΔIC); lag-2(0); arIs51[cdh-3p::GFP]* were observed to have a single AC, indicating that the AC/VU decision was normal.

Loss of maternally-provided APX-1 from the P2 blastomere causes an excess of pharyngeal cells that is evident in later-stage embryos [16], readily assayed by *tbx-38p::mCherry-Histone* expression [18–20]. We depleted *epn-1* in four-cell embryos by feeding parents bacteria expressing double-stranded RNA, an efficient way to deplete maternally-provided transcripts [21]. To ensure efficient depletion had occurred, we created a GFP-tagged knock-in allele of *epn-1* (see STAR Methods). EPN-1::GFP is readily visualized in all four blastomeres at the four-cell stage of embryogenesis, and RNAi efficiently reduces EPN-1::GFP to undetectable levels at the 4-cell stage (Figure 1B, left). We observed that all *epn-1(RNAi)* embryos had a wild-type number of mCherry-Histone-labeled cells, indicating that loss of *epn-1* activity does not prevent APX-1 activation of GLP-1 in the ABp embryonic blastomere (Figure 1B, right).

As a complement to removing EPN-1, we tested the signaling activities of forms of both LAG-1 and APX-1 that were mutated or truncated to block their ability to interact with Epsin per se or with all cytosolic proteins respectively (Figure 1C). Previous studies using high copy number “simple” extrachromosomal arrays suggested that APX-1 could substitute for LAG-2 and that the intracellular domain of both ligands is dispensable for ligand activity when driven by *lag-2* regulatory sequences [22–24]. Such arrays, the only transgene technology available at the time, characteristically cause overexpression and are subject to array-to-array variability, potentially influencing the results or their interpretation. To circumvent these issues, we examined single-copy transgenes inserted into a defined site via CRISPR/Cas9 editing ([25]; STAR Methods), using *lag-2* regulatory sequences to achieve expression of wild-type or mutant LAG-2 or APX-1 at a level expected to be similar to that of the endogenous *lag-2* locus (Figure 1C).

Epsin binds directly to ubiquitinated substrates [26, 27] to target them for Clathrin-mediated endocytosis. We therefore blocked the capacity of either LAG-2 or APX-1 to interact with Epsin by mutating all of the intracellular lysines to arginine [“KtoR”] to prevent ubiquitination. Transgenes encoding either the LAG-2(KtoR) or APX-1(KtoR) fully rescued *lag-2(0)* (Figure 1C), supporting the conclusion that Epsin is not required to provide a pulling force via direct interaction with transmembrane ligands in *C. elegans*. We also tested rescuing transgenes encoding truncated forms of LAG-2 and APX-1 that lack the entire intracellular domain [(ΔIC)] and again observed full rescue of *lag-2(0)* animals, indicating that both ligands can signal normally in the absence of direct interactions with the cytosolic domains of any other endocytic adaptor proteins.

Finally, we also tested the signaling capacity of secreted forms of the LAG-2 and APX-1 ectodomains expressed from single-copy, rescuing transgenes. Although prior studies have reported signaling activity by secreted ligand ectodomains when overexpressed, we did not observe rescue of *lag-2(0)* lethality when expressed from single copy transgenes in the LGI site (Figure 1C). However, when we added a known GPI attachment signal from *vab-2*/Ephrin [28] to the secreted ectodomain of LAG-2, we observed that it restored rescuing activity, albeit with incomplete penetrance (Figure 1C), suggesting that association of the ligand ectodomain with the signaling cell membrane facilitates its signaling activity. We consider the potential significance of this finding in the Discussion.

### The NRRs of LIN-12 and GLP-1 have a lower force threshold for ligand-induced cleavage

The absence of a requirement for Epsin-mediated ligand endocytosis to activate LIN-12 and GLP-1 differs from the absolute requirement observed for Notch signaling in Drosophila [9, 11, 12] and vertebrates [8]. We therefore considered whether this striking difference on the ligand side is due to a corresponding difference on the receptor side. We focused on the NRRs of LIN-12 and GLP-1 because the critical role of ligand is to expose the ADAM protease cleavage site that is buried in the autoinhibited conformation [1, 2]. In particular, we wondered if the LIN-12 and GLP-1 NRRs might be tuned to a lower force threshold, obviating the need for the strong pulling force provided by Epsin-mediated ligand endocytosis. To test this hypothesis, we used chimeric ligand-receptor binding pairs in *Drosophila* to assess the force required to expose the ADAM cleavage site embedded in the NRRs of LIN-12 and GLP-1. This approach was successfully used to establish that the NRR is a bona fide force sensor and that a distinct level of Epsin-mediated mechanical tension across the intercellular ligand/receptor bridge is both necessary and sufficient to induce ADAM-mediated Notch activation *in vivo* [6].

We generated chimeric ligand-receptor pairs in which the ligand/receptor interaction domains of Delta and Notch are replaced by the corresponding interaction domains of vertebrate Follicle Stimulating Hormone (FSH) and FSH Receptor (FSHR), and the native Drosophila Notch NRR is replaced by the NRR of either LIN-12 or GLP-1 (Figure 2x). The chimeric ligands and receptors were then expressed in mutually exclusive sub-populations of cells in the Drosophila wing imaginal disc using Mosaic Analysis by Promoter Swap technology (MAPS; [6]; STAR Methods). Under these conditions, productive signaling that activates the chimeric receptor causes ectopic induction of the Notch target gene *cut* (Figure 2x). We also assayed reliance on Epsin-dependent ligand endocytosis by using mutant forms of the chimeric ligand in which the intracellular domain corresponded to the KtoR, ΔIC, GPI-tethered, and secreted forms of FSH-Dl ligands that were assessed in *C. elegans* (Figure 1C, Figure 2x).

**Figure 2.**
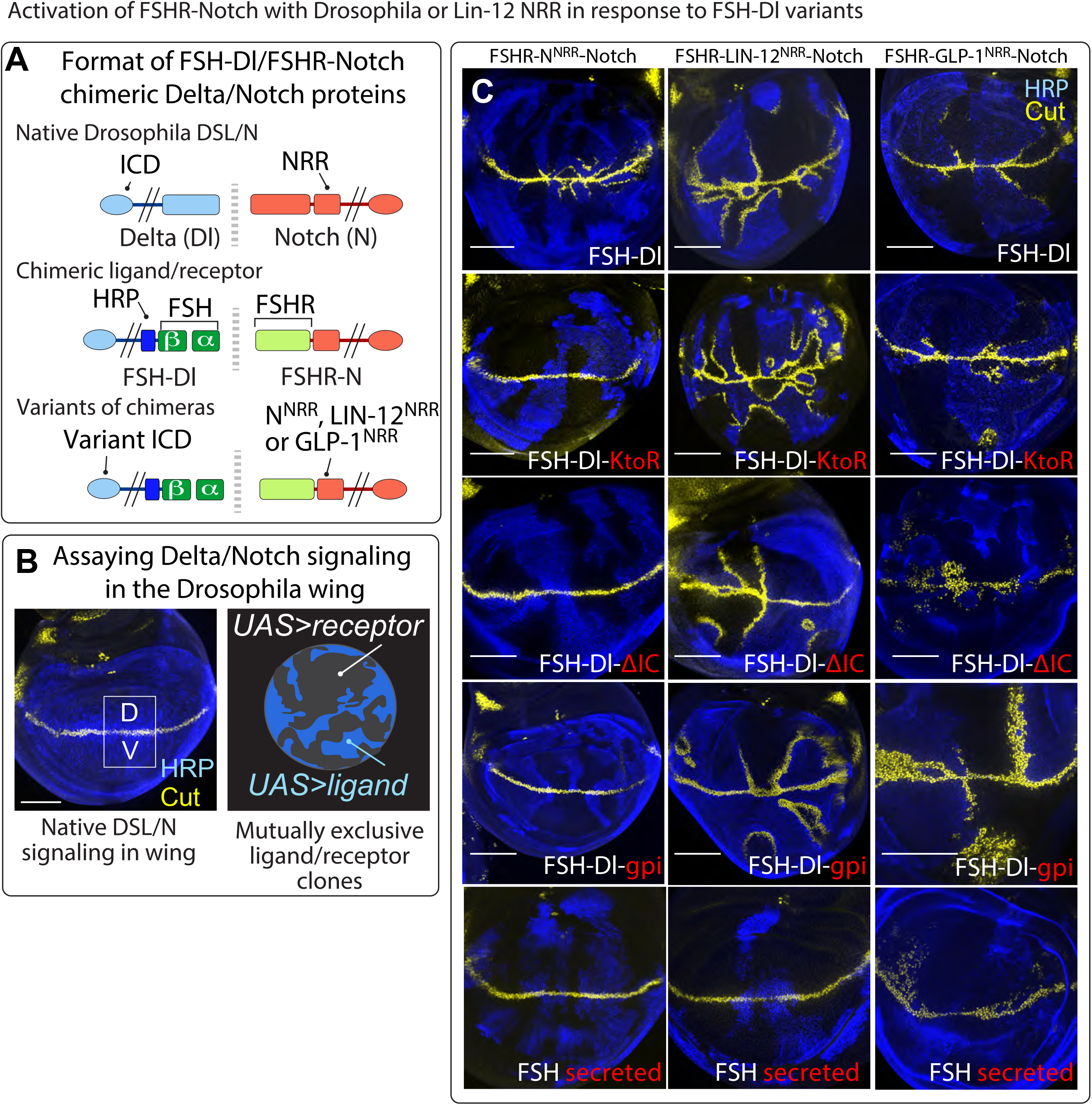
*In vivo* comparison of signal transduction by versions of Notch with the NRR from *Drosophila* Notch, *C. elegans* LIN-12 or *C. elegans* GLP-1. The chimeric version of *Drosophila* Notch is activated only by ligands that enter the Epsin endocytic pathway, whereas the same receptor with the LIN-12 NRR is also activated by ligands excluded from the Epsin endocytic pathway. **A)** Ligand binding regions of Delta/Notch were replaced with functional heterologous binding regions. The Dl extracellular domain (ECD) was replaced by the β subunit of Follicle Stimulating Hormone (FSHβ) and FSHα was expressed to reconstitute the composite FSH ligand (FSH). Reciprocally, the ligand-binding (EGF) portion of the Notch ectodomain was replaced by the FSH receptor ectodomain (FSHR). Further variants were produced by replacing the fly NRR from the canonical receptor (FSHR-N^NRR^-N) with the LIN-12 NRR (to form FSHR-LIN-12^NRR^-N). The ligand was modified by replacing the Delta intracellular domain (ICD) that drives FSH-Dl into the Epsin route with ICDs that exclude the ligand from that route (FSH-Dl-KtoR, -ΔIC or -gpi). **B)** Endogenous Notch activation in the wing primordium results in the Notch target gene *cut* expressed at high levels at the border between the dorsal (D) and ventral (V) compartments (left panel). Ligand/receptor transgenes are expressed under *UAS* control in mutually exclusive subpopulations of receptor or ligand expressing cells distinguished by epitope tagging the ligand, in this case HRP-tagged Dl, stained blue (right panel). **C)** FSHR-N^NRR^-N is activated only by ligands that enter the Epsin endocytic pathway, whereas FSHR-LIN-12^NRR^-N and FSHR-GLP-1^NRR^-N are also activated by ligands excluded from the Epsin endocytic pathway. FSH-Dl can access the Epsin pathway and FSH-Dl expressing cells (blue) induce ectopic Cut (yellow) in adjacent FSHR-N^NRR^-N (top, left column) and FSHR-LIN-12^NRR^-N cells (top, right column). There is no ectopic Cut where FSHR-N^NRR^ -N cells confront ICD variant forms of FSH-Dl that cannot access the Epsin route (left column; FSH-Dl-KtoR, FSH-Dl-ΔIC and FSH-Dl-gpi). In contrast, there is strong ectopic Cut expression in FSHR-LIN-12^NRR^-N cells confronting cells expressing all three variant FSH-Dl forms that do not access the Epsin route (middle column). Similarly, there is strong ectopic Cut expression in FSHR-GLP-1^NRR^-N cells confronting cells expressing the three variant FSH-Dl forms that do not access the Epsin route (right column). Secreted FSH (bottom row) fails to induce Cut expression in neighboring FSHR-N^NRR^-N and FSHR-LIN-12^NRR^-N cells (blue), but does induce Cut expression in neighboring FSHR-GLP-1^NRR^-N cells. Scale bars: 50μm.

In wild-type wing imaginal discs, *cut* expression is tightly restricted to “border” cells flanking the dorso-ventral (D/V) compartment boundary, where DSL/Notch signaling is at a peak [29]. We note that DSL/Notch signaling extends many cell diameters from this boundary, declining as a function of distance, although it is only strong enough to activate Cut expression at the boundary. In MAPS mosaic discs in which FSH-Dl expressing cells abut FSHR-Notch chimeric receptors carrying the Notch NRR (FSHR-N^NRR^-Notch) expressing cells, the strong signaling interfaces extend up to ~10 cell diameters from the D/V boundary (Figure 2x). Activation of FSHR-N^NRR^-Notch depends on the Epsin-mediated pulling force because it cannot be activated by the K-to-R or ΔIC form of ligand, or in the absence of Epsin in the signaling cell [6].

By contrast, in FSHR-LIN-12^NRR^-Notch expressing cells, Cut expression can be induced at distances up to 30 or more cell diameters from the D/V compartment boundary. This markedly more robust response suggests that FSHR-LIN-12^NRR^-Notch is more sensitive to activation by ligand, consistent with the LIN-12 NRR being tuned to a lower threshold than the Notch NRR. This inference is strongly supported by the observation that chimeric FSHR-LIN-12^NRR^-Notch also shows a comparably robust response to both the K-to-R and ΔIC forms of the FSH-Dl ligand, even though they cannot exert the Epsin-dependent pulling force, and also a detectable but somewhat weaker response to the GPI-linked FSH ligand; however, there is no response to the secreted FSH ligand. In sum, the response of the FSHR-LIN-12^NRR^-Notch receptor in the Drosophila MAPS assay parallels that of the response of the LIN-12 receptor to LAG-2 and APX-1 signaling in *C. elegans*. Thus, we conclude that the Epsin-independence of LIN-12 activation in *C. elegans* can be attributed to its NRR being tuned to a lower force threshold than Drosophila Notch.

We also performed similar MAPS experiments testing the response of the chimeric FSHR-GLP-1^NRR^-Notch receptor. At 25°C, we observed some constitutive transducing activity (Figure S1). This constitutive activity is eliminated at 23°C, where activation of the chimeric FSHR-GLP-1^NRR^-Notch is strictly ligand-dependent, allowing us to examine the response to different ligand forms as above. With one exception, the results obtained for the chimeric FSHR-GLP-1^NRR^-Notch receptor at 23°C were similar to those obtained for FSHR-LIN-12^NRR^-Notch at 25°C. The one exception is that FSHR-GLP-1^NRR^-Notch displays some degree of activation by the secreted FSH ligand, whereas FSHR-LIN-12^NRR^-Notch did not, even at 25°C. This exception is consistent with activation of the GLP-1 NRR at a lower force threshold than the LIN-12 NRR.

### Evidence that the “Leucine plug” contributes to tuning the force sensitivity of the NRR

A striking feature of the structure of human Notch NRRs is a “Leucine plug,” a Leu moiety whose side chain is in position to block access to the cleavage site [1, 2, 5]. This residue is highly conserved in Drosophila Notch and vertebrate Notch proteins, but is absent in LIN-12 and GLP-1, even though the surrounding context is conserved. The correlation between the presence of this moiety and the requirement for an Epsin-dependent “pull” to induce the ADAM cleavage of the NRR in most systems led us to test whether the absence of this structural element in LIN-12 and GLP-1 might be responsible for the Epsin-independence of DSL/Notch signaling in C. elegans.

For both the LIN-12 or GLP-1 NRRs, the conserved Leucine thought to constitute the plug is simply absent. Remarkably, when we introduced a Leucine at this position, we found that the modified FSHR-LIN-12^NRR^-Notch and FSHR-GLP-1^NRR^-Notch receptors were now dependent on Epsin-mediated ligand endocytosis (Figure 3). Both modified receptors respond robustly to the wild-type FSH-Dl ligand. In contrast, the response of the plug modified receptors to either the KtoR or ΔIC froms of FSH-Dl is restricted to within a few cell diameters of the DV border. These observations suggest that introducing the leucine plug has increased the force threshold necessary to induce ligand-induced cleavage and activation.

**Figure 3.**
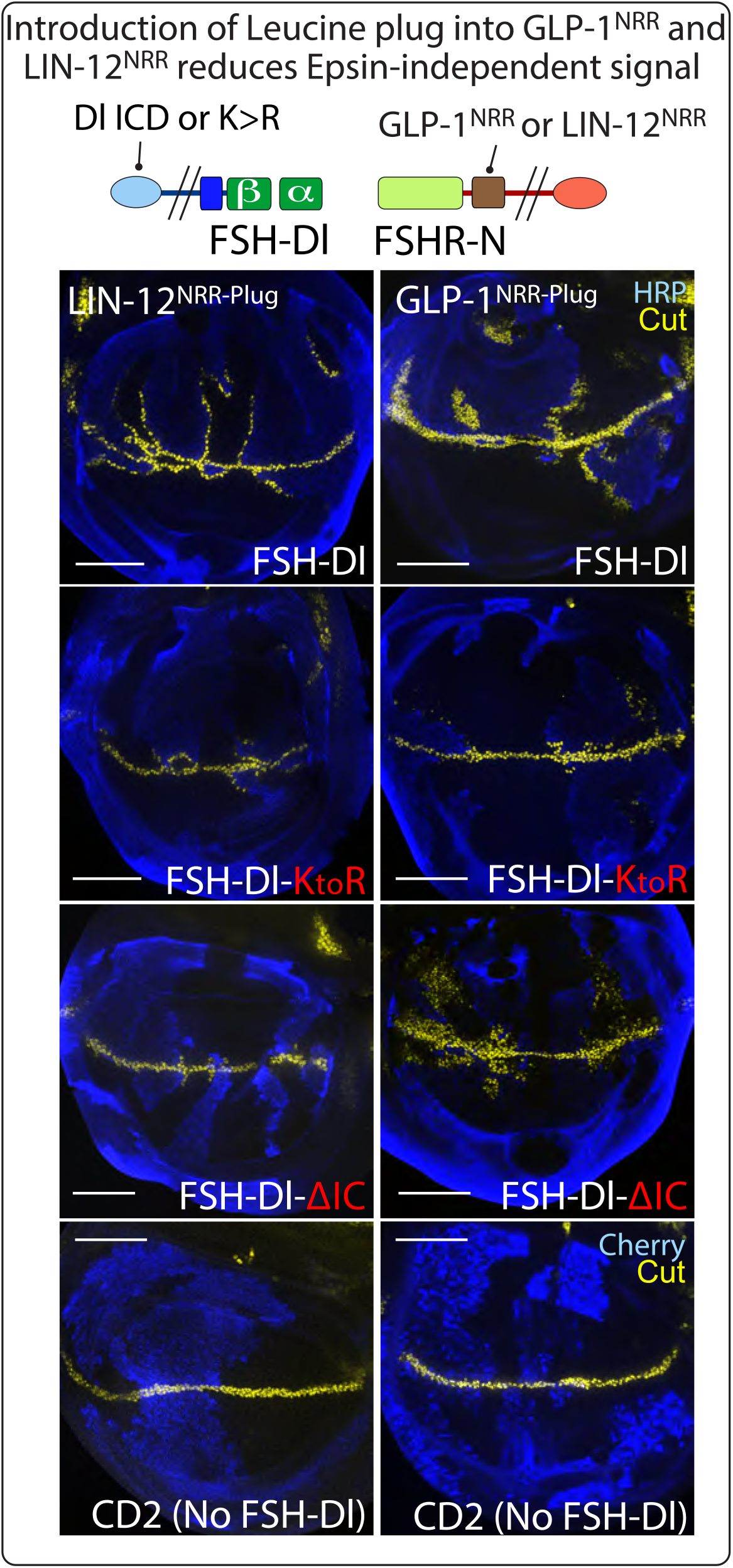
FSHR-N receptors with LIN-12 and GLP-1 NRR variants that include a ‘leucine plug’ show a reduced response to FSH-Dl ligand that is internalized via non-Epsin routes. The NRR of FSHR-LIN-12^NRR^-N and FSHR-GLP-1^NRR^-N was modified by the introduction of a single leucine residue at a conserved position to form FSHR-LIN-12^NRR-Plug^-N (left column) and FSHR-GLP-1^NRR-Plug^-N (right column). Each NRR-Plug variant induces strong Cut expression when confronted with cells expressing FSH-Dl (top row, blue), with response detectable ~30 cell diameters from the DV border. This response is equivalent to that observed for the unmodified FSHR-LIN-12^NRR^-N and FSHR-GLP-1^NRR^-N (see Figure 2). However, both FSHR-LIN-12^NRR-Plug^-N and FSHR-GLP-1^NRR-Plug^-N induce ectopic Cut production that is confined to within ~10 cell diameters of the DV border when confronted with the ligands FSH-Dl-KtoR (second row) and FSH-Dl-ΔIC (third row) that are excluded from internalization via Epsin-mediated endocytosis. This is in contrast to the unmodified FSHR-LIN-12^NRR^-N and FSHR-GLP-1^NRR^-N receptors that show strong activation in response to FSH-Dl-KtoR and FSH-Dl-ΔIC. Scale bars: 50μm.

## Discussion

Notch signal transduction is initiated by binding of ligand expressed on the signaling cell to receptor on the receiving cell, leading to proteolytic cleavage of the Notch ectodomain by an ADAM protease in the receiving cell [3, 4]. In the absence of ligand, a Negative Regulatory Region (NRR) in the Notch ectodomain adopts an autoinhibited conformation, masking the ADAM cleavage site [1, 2]. In Drosophila and vertebrates, the endocytic adaptor Epsin generates a “pulling force” by recruiting the ligand to the Clathrin-mediated endocytic machinery, exposing the cleavage site in the NRR. When ligands bind Notch but do not enter the Epsin pathway, they fail to activate the receptor because they cannot exert sufficient force [6]. Thus, Epsin is an essential core component of the Notch signaling system in Drosophila and vertebrates, such that loss of Epsin results in phenotypes associated with the absence of Notch [8–12].

By contrast, here we have shown that in *C. elegans*, Epsin is not required for Notch signaling and that the intracellular domains of transmembrane ligands do not interact with alternative intracellular machinery in the signaling cell to generate a pulling force. We have accounted for these differences by showing that the ADAM cleavage site embedded in the NRRs of both *C. elegans* Notch proteins, LIN-12 and GLP-1, is exposed at a lower force threshold than Drosophila Notch. Because we were able to recapitulate ligand-dependent activation in the Drosophila assay using just the NRRs from the *C. elegans* Notch proteins in chimeric receptors, we suggest that their conformational change is induced by a pulling force in the signaling cell father than by an allosteric effect mediated by the normal interaction between the native ligand and receptor, even though the threshold for activation is lower (Figure 4).

**Figure 4.**
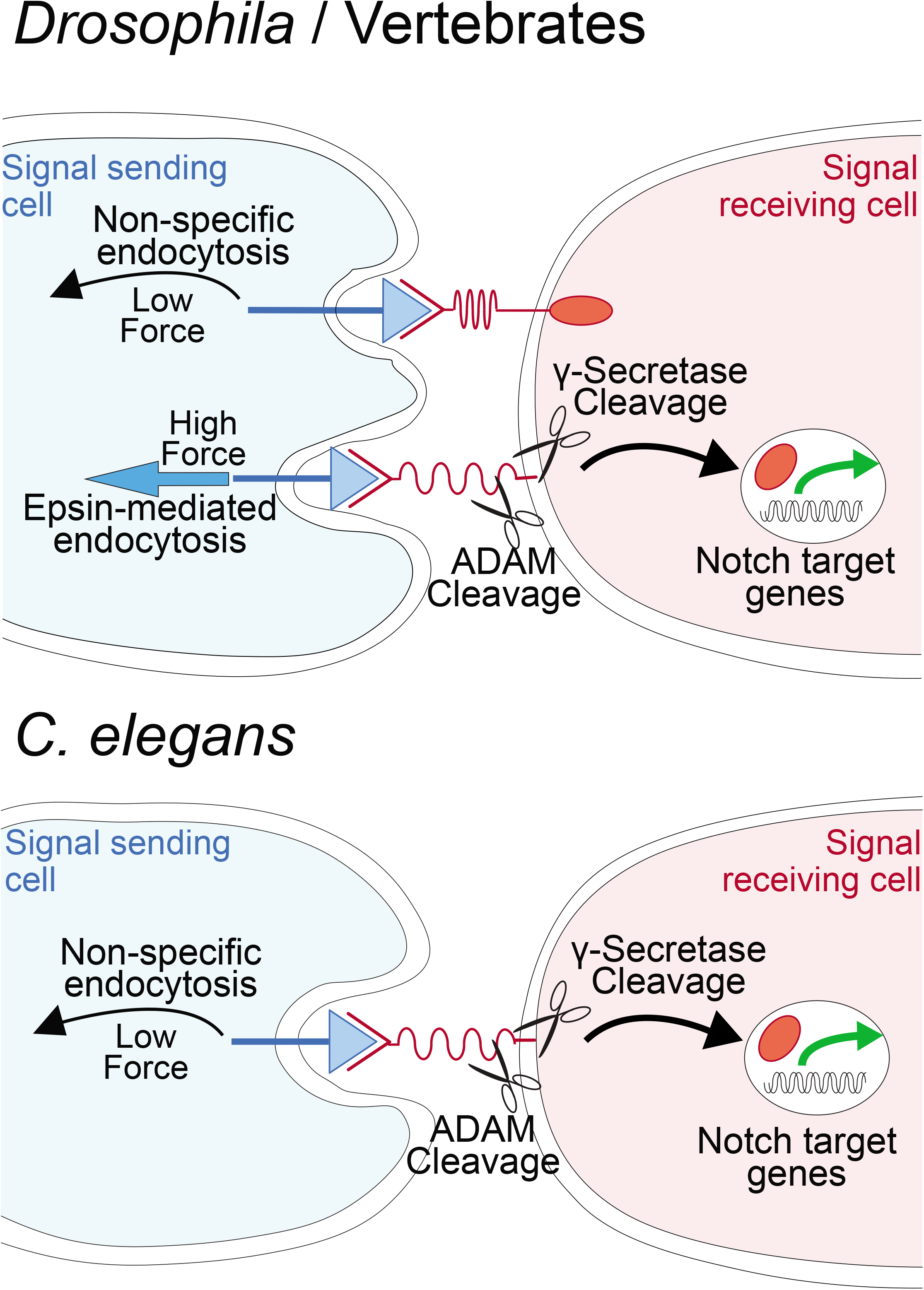
The C. elegans Notch proteins, LIN-12 and GLP-1, are tuned to lower force activation thresholds than Drosophila Notch. In vertebrates and in *Drosophila* (top), there is a strict requirement for Epsin-mediated ligand endocytosis to generate sufficient force to induce the S2 cleavage (scissors) that initiates the activation of the Notch receptor. Without Epsin-mediated endocytosis, ligand and Notch receptor can engage [6], but activating force is not generated, despite ligand internalization via non-Epsin mediated routes. In *C. elegans*, Epsin-dependent endocytosis of the ligand is dispensable for activation as both routes of endocytosis are capable of exerting sufficient force for receptor activation.

What could provide the pulling force if the ligand intracellular domain is dispensable? Expression of the secreted ectodomains of LAG-2 or APX-1 at physiological levels was unable to provide sufficient ligand activity to rescue the lethality of a *lag-2* null allele, a stringent test of ligand function; however, addition of a GPI attachment signal to the secreted ectodomain of LAG-2 significantly restored rescuing activity, indicating that association with the signaling cell membrane potentiates ligand activity. On its own, this observation does not distinguish whether GPI linkage potentiates activity by fostering membrane association per se or by preventing diffusion of the ligand, which would reduce its local concentration. However, the results of the force sensor assay in Drosophila lead us to favor the view that membrane association per se potentiates ligand activity, and raises the possibility that nonspecific endocytic processes such as macropinocytosis or micropinocytosis are sufficient to generate a pulling force (Figure 4). Interestingly, although these endocytic processes are nonspecific, they can be regulated [30, 31], further raising the possibility that signaling by DSL proteins in *C. elegans* may be modulated at this level. It is also possible that the rigidity of the membrane alone may generate sufficient mechanical tension across the ligand-receptor bridge, as immobilizing ligands on a surface such as a culture dish allows for activation of Notch *ex vivo* whereas soluble ligands do not [32–34]. The potentiation of ligand activity by membrane association also suggests that natural secreted ligands may have an as-yet unidentified partner that provides a membrane anchor.

We also changed the tuning of LIN-12 by adding a Leucine moiety that is present in Drosophila and vertebrate Notch proteins, but is naturally absent in *C. elegans* Notch proteins, thereby creating an altered LIN-12 NRR that then becomes dependent on the Epsin-mediated pulling force. This observation supports the view that removing a “Leucine plug” by mechanical force [1,2, 5] is a key step in ligand-induced activation of Drosophila and vertebrate Notch, and further suggests that the Epsin-mediated pulling force is specifically required to accomplish this step.

We note that the intracellular domains of LAG-2 and APX-1 are highly conserved among *Caenorhabditis* species (but not orthologs in other nematode genera) even though they appear to be dispensable for the cell fate specification events we examined. Perhaps in *C. elegans* it calibrates the level of signaling by regulating the production, trafficking, or stability of DSL proteins, and/or plays a role in increasing ligand-dependent force in a specific cell context or physiological condition that we have not assayed here. In this regard, we note that RNAi-mediated reduction of *epn-1* activity reduces the extent of the germline stem cell zone, consistent with a modest effect on DSL signaling to the germline [10], for which both LAG-2 and APX-1 serve redundantly as ligands [35]. *epn-1* activity in the germline niche may contribute to robust ligand-induced activation of GLP-1, and hence more germline stem cells, even if *epn-1* is not strictly required for fertility. Indeed, regulation via the intracellular domain of DSL ligands may contribute to the modulation of ligand signaling from the niche to the germline stem cells that occurs during development and in response to environmental conditions [36]. Alternatively, reducing Epsin (or *mib-1*, the ortholog of a ubiquitin ligase that target DSL proteins in vertebrates [37]) may alter membrane homeostasis so as to affect ligand or GLP-1 activity indirectly. We note also that there is evidence in Drosophila and vertebrates that the relationship between intracellular domain ubiquitylation and ligand activity is complex [6, 38, 39], and may go beyond the obligate requirement for an Epsin-mediated pulling force, e.g. in the release of cis-inhibitory interactions between Delta and Notch that may also calibrate the level of signaling [40–43].

The Notch signaling system is present in animals ranging from sponges to humans [44, 45]. Our results suggest that there is evolutionary plasticity in the tuning of the ligand-receptor interaction, although we cannot yet distinguish whether the lower force tuning operative in *C. elegans* represents an ancestral state, with subsequent co-option of the Clathrin pathway as the major mechanism for activating Notch in higher animals, or loss of a more ancient Clathrin-mediated mechanism in an ancestor to *C. elegans*. The Drosophila force-sensing assay utilized here offers one approach to future exploration of this question.

## Acknowledgements

We gratefully acknowledge advice and assistance for *C. elegans* work from Julie Canman and Sophia Hirsch, and Michelle Attner; Nikola Danev for assistance with making some of the Drosophila constructs; and Gleniza Gomez and Chunyao Tao for technical assistance. We also thank Zheng Zhou, Sophia Hirsch and Julie Canman for providing strains as credited herein; some strains were provided by the CGC, which is funded by NIH Office of Research Infrastructure Programs (P40 OD010440). Research reported in this publication was supported by the Institute of General Medicine of the National Institutes of Health under award numbers R35 GM131746 (to I.G.) and R35 GM127141 (to G.S.). The content is solely the responsibility of the authors and does not necessarily represent the official views of the National Institutes of Health.

## STAR Methods

Further information and requests for resources and reagents should be directed to and will be fulfilled by the Lead Contact, Iva Greenwald.

### Experimental Model and Subject Details

#### *C. elegans* strains

Strain names and full genotypes are listed in Key Resources Table. *Caenorhabditis elegans* strains N2 (wild type) and GS9526 *[lag-2(q411)/tmC16[unc-60(tmIs1237)]* served as the recipients for microinjection to obtain transgenes. *lag-2(q411)* is homozygous lethal and is a likely molecular null allele as it is caused by an early stop codon [24]; the balancer *tmC16* is described in [46] and was obtained from Caenorhabditis Genetics Center. All strains were maintained at 25°C.

Strain ZH1800 *epn-1(en47); enEx867[epn-1::GFP]* [14] was kindly provided by Zheng Zhou.

Strain JCC596 [19], kindly provided by Sophia Hirsch and Julie Canman, includes *zuls178*, which marks all nuclei with HIS-72::GFP, and *stIs10138*, which marks pharyngeal cells with mCherry:HistoneH1 and can be used assess GLP-1/Notch activity at the four-cell stage: it is expressed in approximately 25% of the cells in late-stage embryos when GLP-1 has been activated in ABp, and in approximately 50% of cells when Notch activation in ABp has not occurred.

Two transgene markers were used to facilitate scoring cell fate in mutant larvae: *arIs51[cdh-3p::gfp]* marks the AC [47] and *arIs164[glt-3p::Venus]* marks the excretory cell [48].

Transgenes generated during the course of this study are described in the Method Details below. Strains containing these transgenes analyzed herein are described in the Key Resources Table.

#### *Drosophila* stocks

Stocks used for the analysis herein are described in the Key Resources Table.

### Method Details

#### *C. elegans* Method Details

##### Plasmids and transgenes for *lag-2(0)* rescue

Single-copy insertion transgenes (*arSi*) were generated at the *ttTi4348* locus on LGI using CRISPR-Cas9 by injection into strain N2 with pAP82 at 50 ng/ul, pGH8 at 10 ng/ul, pCFJ90 at 5 ng/ul, and plasmid at 10 ng/ul as described (Pani and Goldstein, 2018).

Plasmids were made by cloning the desired insert into pWZ111 backbone [49] using Gibson Assembly, with pWZ111 digested by SpeI and AvrII. All inserts were generated by PCR with AccuprimeFx. pWZ111 contains the homology arms used for insertion of transgene at LGI site *ttTi4348*, and a Self-Excising Cassette containing *sqt-1p::SQT-1* and *rps-0p::hygG* flanked by *loxN* sites, and a *hsp::Cre*. Unless heat-shocked, any transgene generated from such pWZ111 backbone would result in Roller and hygromycin-resistant animals.

##### Single copy insertion transgenes for *lag-2(0)* rescue

*arSi48[lag-2p::lag-2(+)]* was generated from pJC70, which contains the *lag-2* gDNA sequence. It contains a 3.3kb 5’ upstream sequence of *lag-2* as promoter, driving the expression of *lag-2* 1.5 kb genomic DNA containing all *lag-2* exons and introns, and an additional 1 kb of *lag-2* 3’ UTR.

*arSi49[lag-2p::lag-2(KtoR)]* is a single copy insertion on the LGI site generated from pJC71, with all the intracellular lysines (10 lysines after aa307) mutated to arginine.

*arSi89[lag-2p::lag-2(ΔIC)]* is a single copy insertion on the LGI site generated from pJC106, with a stop codon replacing a fragment from aa311 to aa 403, which results in a protein truncated three amino acids after the transmembrane domain; the 3aa after TM were mutated from KYK to RYR to retain a positively-charged internal sequence to keep the protein anchored in the membrane.

*arSi88[lag-2p::apx-1(FL)]* is a single copy insertion on the LGI site generated from pJC100. It contains the 3.3kb *lag-2* promoter, the *apx-1* cDNA, and the 1kb 3’ UTR of *lag-2. arSi90[lag-2p::apx-1(KtoR)]* is a single copy insertion on the LGI site generated from pJC120 with all the intracellular lysines (4 lysines) mutated to arginine.

*arSi91[lag-2p::apx-1(ΔIC)]* is a single copy insertion on the LGI site generated from pJC121, with a stop codon replacing a fragment from 417aa to 516aa, which results in a protein truncated at 6aa after the transmembrane domain. The 6aa after TM are mutated from SFSKWK to SFSRWR.

*arSi95[lag-2p::lag-2(secreted)-GPI]* is a single copy insertion on the LGI site generated from pJC119. *lag-2(secreted)* is based on the design of the same transgene from Fitzgerald and Greenwald, 1994, with the 20aa GPI attachment sequence from *vab-2*/Ephrin [28] added to the C-terminus.

##### Assessment of *lag-2(0)* rescue

To test the single copy insertion *arSi* transgenes for rescue of *lag-2(0)* lethality, we first generated a balanced intermediate strain *arSi/oxTi559; lag-2(q411)/tmC16*. Transgenes were considered to be rescuing if the strain produced surviving progeny lacking *tmC16*, meaning they are homozygous for *lag-2(q411)*. For transgenes that can rescue *lag-2(0)*, we generated homozygous *arSi; lag-2(q411)* strains and assayed the penetrance of rescue by picking 10-15 egg-laying adult P0s onto plates seeded with OP50, removing them after 3 hours of egg-laying, and then scoring progeny 24 hours later for developmental arrest. All experiments were performed at 25. The genotype of all strains was confirmed by sequencing.

##### Endogenous tagging of *epn-1* and *epn-1/RNAI*)

*epn-1(RNAi)* was used to evaluate a potential role in APX-1 signaling in four-cell embryos. The efficacy was assessed by examining depletion of GFP-tagged endogenous EPN-1, using the protocol described by [50]. We note that this allele has a small in-frame deletion that removes two amino acids, P447 and I448 as well as the ZF1 degron, but has the same GFP expression pattern as EPN-1 ::GFP transgenes and no lethality or any other overt defects suggesting that *epn-1* activity is not significantly compromised.

Strain GS9281, genotype *zuIs178 [his-72::GFP]; stIs10138[tbx-38p::H1-mCherry]; epn-1(ar641)*, was used for *epn-1* RNAi. L4 P0s were placed on RNAi plates containing HT115 bacteria expressing dsRNA specific for *epn-1, lacZ* (negative control) and *apx-1* (positive control) at 25°C as per [51]. In the experiment shown in Figure 1B, 4-cell stage embryos were obtained by dissecting adult P0s after 24 hrs in in 10mM levamisole on a glass coverslip. Embryos were then placed on agarose pads on glass slides and were scored for loss of *epn-1::zf1::GFP* expression, and the glass slides were then placed in 25°C incubator. The same embryos with *epn-1::zf1::GFP* knockdown were then scored for expression of *tbx-38p::H1-mCherry* 2 hours later at ~150 cell stage. A similar RNAi experiment without scoring individuals for EPN-1::GFP depletion was performed using the same RNAi treatment and described in the legend to Figure 1B.

EPN-1::ZF1::GFP knockdown was scored by collecting Z-stacks of GFP fluorescence at 320ms exposure time with a Zeiss spinning disk confocal dual camera system at 40x. Scoring of 150 cell stage embryos was done by collecting Z stacks of GFP (488 nm) and DsRed (561 nm) laser at 170ms and 1500ms exposure time respectively, with a Zeiss spinning disk confocal dual camera system at 40x magnification.

The number of mCherry positive nuclei and GFP nuclei were manually counted using ImageJ with the Cell Counter plugin. Graph was created using PRISM.

##### Scoring of *arIs164* in *epn-1(0)*

To score *arIs164[glt-3p::Venus]* in *epn-1(0)*, 10-15 adults of *epn-1(0); enEx[epn-1::GFP]* were isolated onto an OP50 plate to lay eggs for 3 hours and later removed. 24 hours later, progeny from *epn-1(0); enEx[epn-1::GFP]* were picked and imaged on the GFP channel at 50ms exposure time at a Zeiss Axio Imager D1 microscope with an AxioCam MRm at 40x. *epn-1(0)* progeny were identified by the lack of EPN-1::GFP, which is ubiquitously expressed throughout the animal. Low exposure time was used to show presence of EPN-1::GFP, since arIs164 is very bright.

To score *arIs164* in *lag-2(0), lag-2(0)* progeny from *lag-2(0); arEx2511[lag-2(+)]* lacking a green pharynx were picked and imaged on the GFP channel at 700ms exposure time on a Zeiss Axio Imager D1 microscope with an AxioCam MRm at 40x. Same was done for the control *lag-2(0); arEx2511[lag-2(+)]*, except images were taken at 50ms exposure time to show presence of green pharynx, since arIs164 is very bright. We note that *arEx2511[lag-2(+)]* was generated by injection into *lag-2(q420ts)* with PvuI digested pJC95 at 3ng/ul, ScaI digested pCW2.1 (*ceh-22p::GFP)* at 2ng/ul, and PvuII digested N2 genomic DNA at 50ng/ul and then crossed into the background of the molecular null allele *lag-2(q411)* for phenotypic analysis.

##### Statistical Analysis: *C. elegans*

When comparing two *C. elegans* genotypes for the frequency of two outcomes, a two-tailed 2×2 Fisher’s exact test was used. Differences were considered significant if the p-value is less than or equal to 0.05 for the single-embryo experiment (Figure 1B), and less than or equal to 0.01 for the bulk experiment (Figure 1B legend).

##### *Drosophila* Method Details

###### *Drosophila* Transgenes

All ligand and receptor coding sequences, with the exception of FSHα, were inserted into a modified form of *pUAST-attB* (www.flyc31.org) that contains a single Flp Recombinase Target (FRT, ‘>‘) positioned between the UAS promoter and the coding sequence, and the resulting *UAS>ligand* and *UAS>receptor* transgenes were introduced at a single genomic docking site, *attP-86Fb* located on the right arm of the third chromosome, oriented so that the promoter is centromere proximal to the coding sequence. A “no promoter” (*Ø*) element encoding the transcriptional terminating 3’UTR of the *hsp70* gene occupied the promoter region in *Ø>ligand* transgenes. A single *UAS.FSHα*, transgene inserted by conventional P-element mediated transformation onto the X chromosome was used in all experiments.

The various chimeric forms of Dl are described in more detail in [6]. Briefly, the native extracellular domain of Dl was replaced in its entirety by the β subunit of human Follicle Stimulation Hormone (FSHβ) [52] and the HRP tag was inserted immediately downstream of the heterologous ligand domain, immediately upstream of the Dl transmembrane domain. All versions of FSH-Dl in this work carried an HRP tag and for simplicity this is omitted from their designation, except where necessary to ensure clarity.

The Dl intracellular domain was deleted (ΔIC), mutated so that all Lysines were changed to Arginine (K>R), or replaced just after the stop transfer sequence downstream of the transmembrane domain by wildtype or mutant versions of (i) a small heterologous peptide containing two Lysines that are sufficient to mediate Epsin-dependent endocytosis [11, 12], and (ii) six repeats of a KtoR form of the classic Myc epitope tag of which either five (myc) or six (myc^mut^) are mutated to change the LI dipeptide to AI. To reconstitute the composite FSHα/FSHβ ligand domain, secreted FSHα was co-expressed from the *UAS.FSHα* transgene.

The various chimeric forms of Notch are based on the FSHR-N receptor fully described in [6]. All versions of FSHR-N in this work carried the extracellular Cherry tag and for simplicity this is omitted from their designation, except where necessary to ensure clarity. Briefly the amino-terminal Epidermal Growth Factor (EGF) Repeat containing portion of the native extracellular domain of N was replaced by the ectodomain of FSHR preceded by the signal-peptide from FSH. The extracellular domain was tagged by the insertion of Cherry just upstream of the juxtamembrane NRR or A2 domain; the intracellular domain was tagged by a centrally located insertion of a V5 tag [6]. The extracellular, juxta-membrane NRR was replaced by various C. elegans force sensor domains (Figure S1). In each case, the domain was inserted using an AvrII site and a Not1 site.

Complete DNA sequences are available on request.

##### Analysis at the interface of ligand and receptor cells

Signaling between dedicated ligand and receptor cells was analyzed using Mosaic Analysis by Promoter Swap (MAPS), as fully described in [6]. In essence, mitotic recombination across the FRTs in cells transheterozygous for *UAS>* and *Ø>* transgenes is induced in the presence of a *nub.Gal4* driver that acts in the developing wing. This subdivides the prospective wing into mutually exclusive ligand and receptor expressing subpopulations, allowing signaling to be monitored by assaying the ectopic induction of the Notch target gene *cut* wherever the two subpopulations abut.

To induce mosaics by promoter swap, first or second instar larvae of the appropriate genotype were heat shocked at 36°C for one hour and wing discs from mature third instar larvae were dissected and processed as described above.

##### Quantification and Statistical Analysis: Drosophila

In all experiments, most if not all of the imaginal wing discs contained several mutually exclusive subpopulations of ligand and receptor expressing cells within each wing primordium. In all cases, the images shown in the Figures are representative, and the outcome of the experiments qualitatively apparent (*e.g*., in showing the presence and extent of ectopic Cut expression).

For simple MAPS experiments in which mutually exclusive ligand and receptor expressing subpopulations were generated in otherwise wild type discs, at least 20 discs were scored.

