## Supplemental Figure 1 for "The C. elegans Notch proteins LIN-12 and GLP-1 are tuned to lower force thresholds for activation than Drosophila Notch"

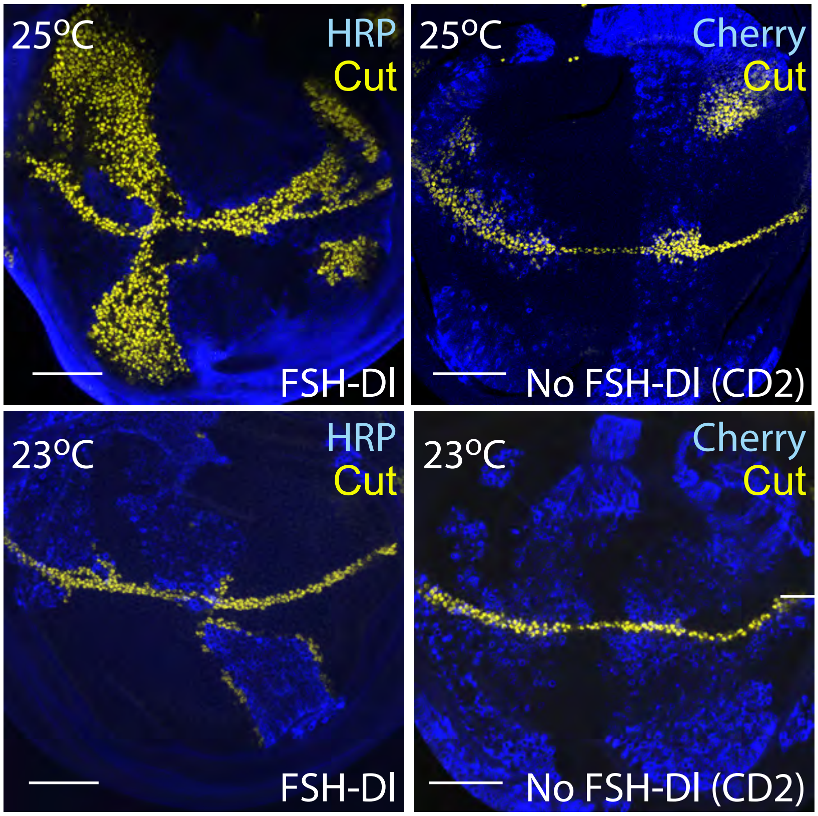


**Supplemental Figure 1.** FSHR-GLP-$1^{NRR}$-N shows some constitutive activity at 25$^{O}$C, but strictly ligand-dependent activation at 23$^{O}$C. At 25$^{O}$C (top row) FSHR-GLP-$1^{NRR}$-N expressing cells produce Cut (yellow) even when located far from FSH-Dl cells (blue, top left) or when CD2, rather than FSH-Dl, is expressed in neighboring clones. In contrast, at 23$^{O}$C (bottom row) FSHR-GLP-$1^{NRR}$-N expressing cells produce Cut only when adjacent to cells expressing FSH-Dl (left) and no ectopic Cut is produced when the ligand is absent from neighboring clones (right). Scale bars: 50µm.
